# Identification of genomic regions affecting production traits in pigs divergently selected for feed efficiency

**DOI:** 10.1101/2020.10.28.358564

**Authors:** Emilie Delpuech, Amir Aliakbari, Yann Labrune, Katia Fève, Yvon Billon, Hélène Gilbert, Juliette Riquet

**Author notes:** Corresponding author: JR. Email addresses: ED.

## Abstract

**Background:** Feed efficiency is a major driver of the sustainability of pig production systems. Understanding biological mechanisms underlying these agronomic traits is an important issue whether for environment and farms economy. This study aimed at identifying genomic regions affecting residual feed intake (RFI) and other production traits in two pig lines divergently selected for RFI during 9 generations (LRFI, low RFI; HRFI, high RFI).

**Results:** We built a whole dataset of 570,447 single nucleotide polymorphisms (SNPs) in 2,426 pigs with records for 24 production traits after both imputation and prediction of genotypes using pedigree information. Genome-wide association studies (GWAS) were performed including both lines (Global-GWAS) or each line independently (LRFI-GWAS and HRFI-GWAS). A total of 54 chromosomic regions were detected with the Global-GWAS, whereas 37 and 61 regions were detected in LRFI-GWAS and HRFI-GWAS, respectively. Among those, only 15 regions were shared between at least two analyses, and only one was common between the three GWAS but affecting different traits. Among the 12 QTL detected for RFI, some were close to QTL detected for meat quality traits and 9 pinpointed novel genomic regions for some harbored candidate genes involved in cell proliferation and differentiation processes of gastrointestinal tissues or lipid metabolism-related signaling pathways. Detection of mostly different QTL regions between the three designs suggests the strong impact of the dataset on the detection power, which could be due to the changes of allelic frequencies during the line selection.

**Conclusions:** Besides efficiently detecting known and new QTL regions for feed efficiency, the combination of GWAS carried out per line or simultaneously using all individuals highlighted the identification of chromosomic regions under selection that affect various production traits.

## Background

Feed efficiency is a major driver of the sustainability of pig production systems. It represents from 50 to 83 % of cost productions depending on the countries and systems [1]. The feed efficiency is also a principal lever to reduce the environmental footprints of the production [2]. The cost of feeding in pig production is usually measured by the computation of feed conversion ratio (FCR). Indeed, FCR is a ratio between two traits of interest for most breeding schemes (feed intake and growth rate), and incorporating it in selection indexes makes it difficult to accurately anticipate responses to selection on this trait and the correlated traits [3]. In 1963, Koch et al. [4] proposed residual feed intake (RFI) as an alternative to quantify feed efficiency, to overcome the limits of FCR. The RFI is the difference between individual feed intakes and predicted feed intake for the animal maintenance and production requirements. It is generally computed as a multiple linear regression of daily feed intake on production traits (growth rate and body composition traits in growing animals), and on the average metabolic body weight of the animal during the period, as an indicator of maintenance requirements. As a result, selection for RFI generates limited correlated responses on the other production traits, as shown in several selection experiments in pigs [5, 6], and other species [7]. However, accurate individual feed intake recording for pigs raised in groups is costly, and large efforts are devoted to facilitate the improvement of feed efficiency, by either identifying biomarkers [8, 9] or genomic markers (for instance [10, 11]). Despite these efforts, the difficulty to find quantitative trait loci (QTL) or genomic variant affecting feed efficiency related traits translates in the PigQTLDB statistics [12]: only 394 QTL are listed for feed conversion type of traits, and 350 for feed intake type of traits, whereas more than 2,000 are reported for growth traits, and more than 3,200 for fatness traits (PigQTLDB, access Sept 2020, https://www.animalgenome.org/cgi-bin/QTLdb/SS/index). Genomic information acquired in established divergent lines for the trait of interest can be used to increase the power of detection of genomic variants for lowly heritable or highly polygenic traits, as RFI in pigs [10] and litter traits in rabbits [13].

In this study, we aimed at identifying genomic regions affecting RFI and other production traits in two pig lines divergently selected for RFI during 9 generations [5], by combining an extensive genotyping of all breeding animals of the lines, and the extensive phenotyping of their progeny. GWAS were applied to growth, feed intake and feed efficiency, carcass composition and meat quality traits on the full dataset. Different subsets of the population were used to suggest biological hypotheses for the genetic background of the traits in the two divergent lines, and decipher whether the chromosomic regions affecting RFI differed between lines.

## Methods

### Ethic statement

All pigs were reared in compliance with national regulations and according to procedures approved by the French Veterinary Services at INRA experimental facilities. The care and use of pigs were performed following the guidelines edited by the French Ministries of High Education, Research and Innovation, and of Agriculture and Food (http://ethique.ipbs.fr/sdv/charteexpeanimale.pdf).

### Design

The data were obtained from a divergent selection experiment on RFI carried out at the INRA experimental units GenESI since 2000 (Surgères, France, https://doi.org/10.15454/1.5572415481185847E12), on growing pigs from the French Large-White (LW) population. The selection procedures were described by Gilbert et al. [5]. In brief, the lines were established from 30 matings of LW animals (F0). From these litters, 116 males were tested to select the 6 most efficient (LRFI) and 6 least efficient (HRFI) males as founders of two divergent lines, and about 40 pairs of sibs were randomly assigned to each line. In the following generations, from G1 to G9, 96 males from each line were tested for RFI to select 6 extreme low or high boars depending on the line. In addition, 35 to 40 females were randomly chosen within-line in each generation to produce the next generation. No selection was applied for females. From G1, matings were organized for at least two successive litters. Until G5, the first litter provided boars candidates for selection and future breeding females, and castrated males and females from the second parity were tested to evaluate the direct and correlated responses to selection on major production traits, including carcass composition and meat quality traits. After G5, selection was applied to parity 4 or 5, and responses to selection were measured on pigs born in parity 2 and 3. Hereafter, the breeding animals will be called “breeders” and animals tested for responses to selection will be called “response animals”.

### Phenotypes

In each generation, 48 females and 48 castrated males per line were produced as response animals, and tested individually during the growing-finishing period (∼28 kg to ∼107 kg) for body weight (BW0 at the start of the test and BW1 before slaughter) and daily feed intake (DFI) using a single-place electronic feeder (ACEMA 64; Skiold Acemo, Pontivy, France) to compute average daily gain (ADG) and feed conversion ratio (FCR) during the test period. The dressing percentage (DP) was computed based on weight records of warm carcass at slaughter. Twenty four hours after slaughter, backfat thickness measured on carcass (carcBFT), and the weights of ham (Ham_W), loin (Loin_W), belly (Belly_W), shoulder (Shoulder_W), and backfat (BF_W), following a standardized cut, were recorded on the cold half carcass. The lean meat content (LMCcalc) was evaluated according to the method of Daumas [14]. Meat quality measurements included pH on *adductor femoris* muscle (AD), *semimembranosus* muscle (SM), *gluteus superficialis* muscle (GS), and *longissimus dorsi* muscle (LM), colorimetry L*, a* and b* on GS and *gluteus medius* muscle (GM), and water-holding capacity (WHC) assessed on GS according to the procedure described by Charpentier et al. [15]. Finally, meat quality index (MQI) was calculated from measurements of pH in SM, L* on GS and WHC according to the model proposed by Tribout et al. [16]. RFI was defined as the residual of a multiple linear regression as follows: RFI = DFI – (1.48 × ADG) + (23.2 × LMCcalc) – (99.1 × AMBW), where AMBW is the average metabolic body weight during the test period and is equal to (BW1^1.6^ −BW0^1.6^)/[1.6 (BW1 − BW0)] [17]. Contemporary group, gender and pen size were added as fixed effects in the model, as described by Gilbert et al.[5].

### Genotyping

Genomic DNA was purified from individual biological samples of the sires and dams of all generations using standard protocols. Over time, two different Illumina medium density SNPs chips were used according to the genotyping protocols defined by the supplier (at Technological Center, Genomics and Transcriptomics Platform, CRCT Toulouse). First batch comprising 286 animals was genotyped for 64,232 SNPs using the Porcine SNP60v2 BeadChip (60K SNPs chip), and a second batch of 1,356 animals was genotyped using the Porcine HD Array GGP chip comprising 68,516 SNPs (70K SNPs chip). Genotypes were obtained using the Genome Studio software (V2.0.4) and coded as 0, 1 and 2 corresponding, respectively, to individuals homozygous for the minor allele, heterozygous and homozygous for the major allele. In addition, 32 G0 founders (12 G0 sires, and 20 G0 dams that had most contribution to the subsequent generations) were genotyped with the Affymetrix Axiom Porcine HD Genotyping Array chip (Gentyane Plateform, UMR 1095 INRAE Clermont-Ferrand) consisting of 658,692 SNPs (650K SNPs chip).

For each SNPs panel, quality control was performed using PLINK software (V1.90) [18]: SNPs with a call frequency (CF) < 95% and a minor allele frequency (MAF) < 1% were excluded, and animals with a call rate (CR) <90% were discarded. Unmapped SNPs and SNPs located on sex chromosomes were removed following the Sscrofa11.1 assembly of the reference genome (https://www.ensembl.org/Sus_scrofa/Info/Index)[19].

### Genotypes imputation

Two successive imputations were performed using the FImpute software [20]. A first level of imputation was performed with markers of 60K and 70K SNPs chips, based on 29,957 SNPs in common, to homogenize the medium density genotyping data available for the 1,632 breeders of the lines. This leads to an intermediate dataset of 66,988 SNPs imputed from both medium density (MD) chips (60K and 70K SNPs chips). In a second step, the genotypes of the high density (HD) SNPs chip were imputed for all breeders using the HD SNPs genotypes of the 32 G0 founders. A set of 45,708 SNPs was in common between MD imputed genotypes and HD SNPs chip. A total of 570,447 SNPs distributed over the 18 pig autosomes, was finally available for 1,632 breeding animals.

To evaluate the imputation accuracy, first, five successive batches of 1,000 SNPs were randomly selected among the common SNPs in the 60K and 70K SNPs chips. For each SNPs batch, the genotypes of these SNPs were set as missing for all animals genotyped with the 60K SNPs chip and imputed from the 70K SNPs chip information. Therefore, a total of 5,000 SNPs with real and imputed genotypes were used to compute Pearson correlations for each of the 286 pigs with 60K genotypes. Similarly, five batches of 1,000 SNPs were randomly selected from common markers of both MD SNPs chips, animals genotyped with the 70K SNPs support were re-coded as missing, and Pearson correlations between true and imputed genotypes were computed for the 1,346 animals with 70K SNPs genotypes. Then, to evaluate the imputation quality to the HD, the same strategy of removing successively five batches of 1,000 SNPs from the data was applied using SNPs in common to the three chips. In addition, a leave-one-out approach was applied to the 32 individuals with HD genotypes to evaluate the imputation accuracy.

In addition, a multi-dimensional scaling (MDS) analysis was performed using R software (V.3.6.2, R Core Team 2019) based on a identity-by-state matrix constructed with the PLINK software [21].

### Predicted genotypes in response animals

Response animals did not have genotypes themselves. An average expected genotypes of their parents was computed for each animal from the imputed 650K genotypes. For each marker, each individual was given the average genotype of the parents (0, 0.5, 1, 1.5 or 2), so within a litter, all animals were assigned the same genotypes. Depending on the genotypic class, the obtained genotype was, therefore, an approximation of the real genotype: (*i*) genotypes 0 and 2 were certain, as they resulted from two homozygous parents for the same allele (0×0 ⟶ 0 and 2×2 ⟶ 2), (*ii*) genotypes 0.5 and 1.5 included combinations of a homozygous genotype for one allele and a heterozygous genotype (0×1 ⟶ 0 or 1 and 1×2 ⟶ 1 or 2), and (*iii*) genotype 1 was the most heterogeneous class, with a mixture of true genotypes (0×2 ⟶ 1) and uncertain genotypes (1×1 ⟶ 0 or 1 or 2). Animals with a parent with a missing genotype were excluded from the analysis.

### Genome-Wide Association Studies

GWAS analyses were performed using GEMMA software (version 0.97) [22] on response animals with their own phenotypes and their average genotypes from parents. Phenotypes were adjusted for significant fixed effects and covariates (pen size, herd, sex, and contemporary groups for *in vivo* measurements, slaughter date as fixed effects, and slaughter age as covariate for traits recorded at the abattoir, and slaughter BW as covariate for carcBFT) using linear models as proposed in Aliakbari et al. [23]. The resulting residues were integrated as phenotypes in GEMMA. To account for the structure of the population in the GWAS analyses, a pedigree relationship matrix **A** was computed. Association analyses were performed on the 24 traits available for 2,426 response animals.

The statistical model used to test one marker at a time was **y = x***β* **+ u + ε**, where **y** is the vector of adjusted phenotypes for all individuals; **x** is a vector of genotypes at the tested marker; *β* is the effect of the tested marker; **u** is a vector of random additive genetic effects distributed according to *N*(0, **A**λτ^−1^), with λ the ratio of the additive genetic variance and the residual variance τ^−1^; **ε** is a vector of residuals *N*(0, Iτ^−1^), with **I** the identity matrix. In GEMMA, an efficient exact algorithm is implemented to first estimate *λ*, and next derive 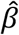 and 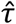for each marker [24].

The distributions of the *p*-values for the GWAS of each trait were checked using quantile-quantile plots (Q-Q plot) and computing regression coefficients of the −log_10_(observed *p*-values) on the −log_10_(expected *p*-values under H_0_). Inflation factors were lower than 1.23 for all analyses, indicating low deviations from the distribution of the test statistic under H0. A correction factor was anyway applied to all analyses to control type-I errors, by dividing each *p*-value by the corresponding inflation factor to avoid the impact of this low deviation.

To account for the nominative type-I error and multiple testing issue, the significance threshold was obtained after a Bonferroni correction as follows:

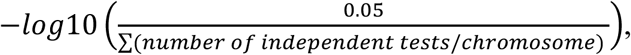

where number of independent tests was computed per chromosome as the number of principal components required to describe 99.5% of the genotypic variability of each chromosome, after a principal component analysis applied to the correlation matrix between genotypes of the SNPs of the considered chromosome (square root (r^2^) of linkage disequilibrium (LD) between each pair of SNPs), as recommended by Gao et al. [25]. The chromosome-wide thresholds obtained were between 3.09 and 3.16, thus a threshold of 3 was used to identify suggestive associations. To determine genome-wide significance thresholds, the number of independent tests for the 18 autosomes was summed to apply a correction at the genome level. This threshold (4.5) was used to identify significant associations.

Three types of populations were considered for GWAS. First, the full dataset combining the two lines was analyzed in a global analysis (thereafter called Global-GWAS). Then, to evaluate if some QTL were segregating in one line only, the analyses were repeated within line (thereafter called Lines-GWAS, or HRFI-GWAS and LRFI-GWAS when only one line was referred too).

To define QTL intervals, for each combination of population and trait, the genome was divided into 1 Mb windows following the Sscrofa11.1 assembly of the reference genome. The 1 Mb windows with at least one SNP with significant *p*-value at 5% genome-wide (-log_10_(*p*-value)≥4.5) were retained, and adjacent windows with significant signals were combined into a single QTL window per trait. In a second step, the adjacent and overlapping significant windows between traits were combined using the same approach as presented above, thus allowing a complete list of QTL regions to be subsequently analyzed. When a QTL region was significant for several traits, for each of them, the most significant marker and the associated allelic substitution effect was retained to tag the QTL (trait x region) for this trait in further analyses – thereafter called SNP-QTL.

The QTL positions were compared to previously mapped QTL in pigs using the pigQTLdb database [12], and QTL significant for RFI trait were screened for functional candidate genes using Ensembl annotation V.101 (August 2020).

### Changes of allelic frequencies of SNP-QTL

The power of detection in GWAS is strongly influenced by the allelic frequencies of the analyzed markers [26]. Within each QTL window, the most significant SNP was considered to examine the changes of allele frequencies with line selection. These SNP-QTL allele frequencies were estimated for the response animal genotypes, i.e. from their average genotypes. To find out how selection affected allele frequencies, and thus power of detection, allele frequencies were computed by adding animals from one generation at a time, starting from G1 individuals alone. Then, the allele frequencies adding G2 response animals were obtained by combining genotypes of G1 and G2 response animals, and so on until G9. The estimated frequencies in G9 (using all the animals from G1 to G9) corresponded to the informativeness of the markers used in the main GWAS by line. A regression of the generation (1 to 9) on the SNP allele frequencies was then applied to test changes of allelic frequencies on cumulative datasets over generations. For each SNP-QTL, the significance of the slope was estimated in each line using a Wald test. An average evolution score was compute for each QTL region (9 generations * (|slope _HRFI_| + |slope _LRFI_|)) when the slope value was different from zero with p < 0.05. To reflect the evolution per trait, an average value over all the QTL regions detected for each trait was computed.

## Results

### Genotype quality control and imputation

True SNPs genotyping data were available for all sires and dams from G0 to G9. The quality control of the genotypes was first carried out for each SNP chip independently. With a CR threshold of 90%, 10 animals genotyped with the 70k SNPs chip and no individual genotyped with the 60K and 650K SNPs chips were discarded (Additional file 1). For the SNPs, 15,114 SNPs from the 60K SNPs chip (5,776 for CF < 95% and 9,125 for MAF < 1%), 11,891 SNPs from the 70K SNPs chip (5,323 for CF < 95% and 6,568 for MAF < 1%), and 99,587 SNPs from the HD SNPs chip (53,735 for CF < 95% and 45,852 for MAF < 1%) were removed. In total, genotypes of 286 animals for 49,118 SNPs for the 60k SNPs chip, genotypes for 1,346 animals for 56,625 SNPs for the 70K SNPs chip, and finally genotypes for 32 animals for 559,105 SNPs for the HD SNPs chip were retained for further analyses (Additional file 2).

To obtain HD genotypes for all parents of the design, two successive runs of imputations were performed. First, the imputation of the missing genotypes on each MD support (60K and 70K SNPs chips) allowed obtaining genotypes for 66,988 SNPs for all sires and dams. The imputation accuracy was on average 0.995 regardless the generation of the imputed individuals (Figures 1a and 1b). A second run of imputation was applied to all breeding animals from the 32 founder individuals genotyped with the HD SNPs chip. The imputation accuracy was also high, with average accuracies around 0.979 (Figure 1c). Some few animals in G0 and G3 had accuracies lower than 0.97. The accuracy estimated via the leave-one-out approach confirmed the values estimated with the correlations, with an average of 0.975. In total, genotypes for 570,447 SNPs were obtained for all parents from G0 to G9.

**Figure 1.**
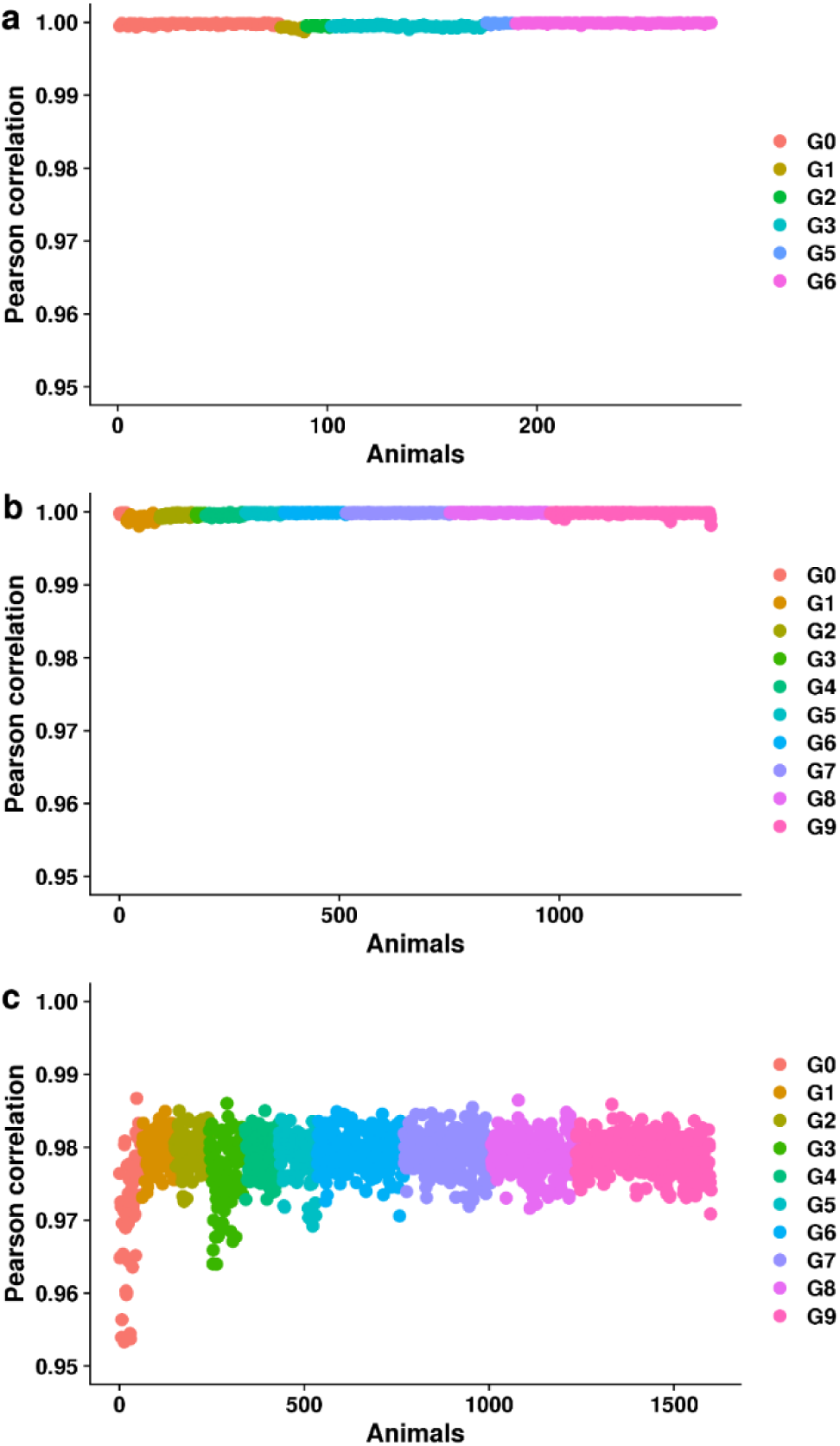
Correlations between true and imputed genotypes for animals genotyped on 60K or 70K SNPs chip. For each analysis, correlations were estimated setting 5,000 SNPs as missing (5 batches of 1,000 SNPs) on one chip among SNPs in common between the two supports used. Animals are sorted and colored by generation. Correlations between true and imputed genotypes (a) for the 286 animals genotyped with the 60K SNPs chip using animals with 70K genotypes as reference population, and (b) for the 1,346 animals genotyped with the 70K SNPs chip using animals with 60K genotypes as reference. (c) Correlations between true and imputed genotypes after imputation to 650K SNPs from the imputed medium density genotypes.

An MDS analysis was performed on the genotypic matrix to represent the changes of genomic content of the lines with generations (Figure 2). The first component corresponded to the dispersion of individuals according to the lines, and the second component corresponded to the successive generations in both lines.

**Figure 2.**
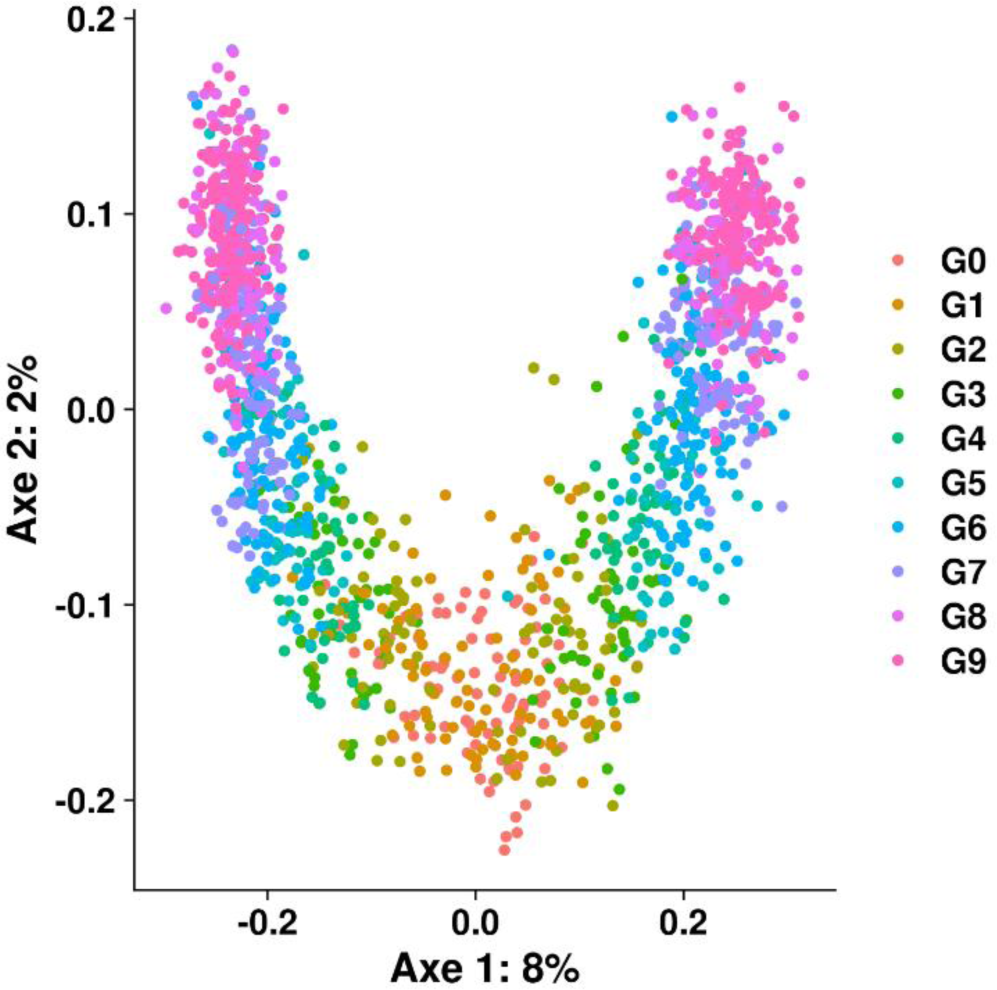
Two first axes of the multidimensional scaling (MDS) analysis, based on the 570,447 genotypes. Points represent individuals (corresponding to all sires and dams of the population, N=1,632) and colors are generations.

### Genome-wide association studies

From the imputed genotypes of all parents, an average genotype was computed for all response animals. Thus genotypes coded 0, 0.5, 1, 1.5 or 2 were available for 2,426 individuals in total. Within a sibling, all individuals shared the same average genotype. On average the size of the siblings was 4.07 (± 2.9).

First, association studies corresponding to Global-GWAS were carried out on all response animals, for each of the 24 traits. A total of 54 regions of 1 Mb (38 regions), 2 Mb (12 regions), or 3 Mb (4 regions) were significant for at least one trait, corresponding to 72 QTL (trait x region). QTL were detected for all 24 traits (Figure 3), the list and characteristics of these QTL is reported in the Additional file 3. Cut weights were the traits with the lower number of QTL (1 to 3 per analysis), except for the weight of backfat (BF_W) in the Global-GWAS (Table 1). Meat quality measurements had the highest number of QTL (up to 7). Thirty regions associated with growth, feed intake, and feed efficiency were detected, including 12 regions associated with RFI and 5 with FCR. For all traits (except Belly_W), at least one QTL was detected in the Global-GWAS.

**Table 1.**
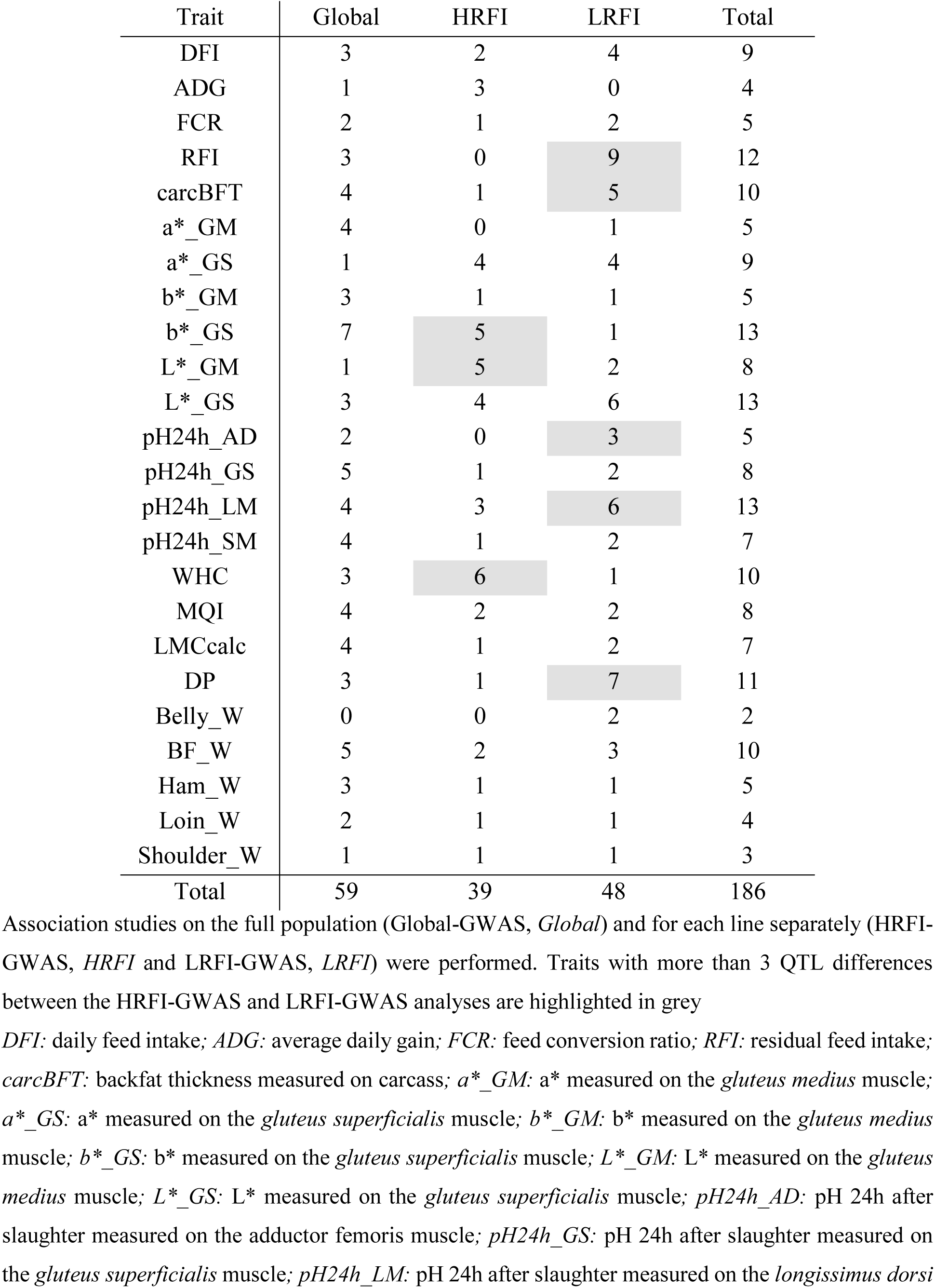
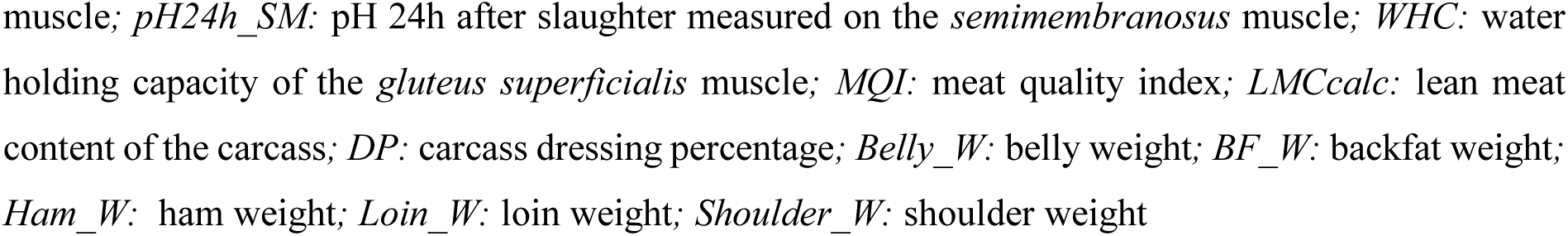
Number of QTL identified for each trait with the 3 groups of association studies.

| Trait | Global | HRFI | LRFI | Total |
| --- | --- | --- | --- | --- |
| DFI | 3 | 2 | 4 | 9 |
| ADG | 1 | 3 | 0 | 4 |
| FCR | 2 | 1 | 2 | 5 |
| RFI | 3 | 0 | 9 | 12 |
| carcBFT | 4 | 1 | 5 | 10 |
| a*_GM | 4 | 0 | 1 | 5 |
| a*_GS | 1 | 4 | 4 | 9 |
| b*_GM | 3 | 1 | 1 | 5 |
| b*_GS | 7 | 5 | 1 | 13 |
| L*_GM | 1 | 5 | 2 | 8 |
| L*_GS | 3 | 4 | 6 | 13 |
| pH24h_AD | 2 | 0 | 3 | 5 |
| pH24h_GS | 5 | 1 | 2 | 8 |
| pH24h_LM | 4 | 3 | 6 | 13 |
| pH24h_SM | 4 | 1 | 2 | 7 |
| WHC | 3 | 6 | 1 | 10 |
| MQI | 4 | 2 | 2 | 8 |
| LMCcalc | 4 | 1 | 2 | 7 |
| DP | 3 | 1 | 7 | 11 |
| Belly_W | 0 | 0 | 2 | 2 |
| BF_W | 5 | 2 | 3 | 10 |
| Ham_W | 3 | 1 | 1 | 5 |
| Loin_W | 2 | 1 | 1 | 4 |
| Shoulder_W | 1 | 1 | 1 | 3 |
| Total | 59 | 39 | 48 | 186 |

Association studies on the full population (Global-GWAS, *Global*) and for each line separately (HRFI-GWAS, *HRFI* and LRFI-GWAS, *LRFI*) were performed. Traits with more than 3 QTL differences between the HRFI-GWAS and LRFI-GWAS analyses are highlighted in grey

**Figure 3.**
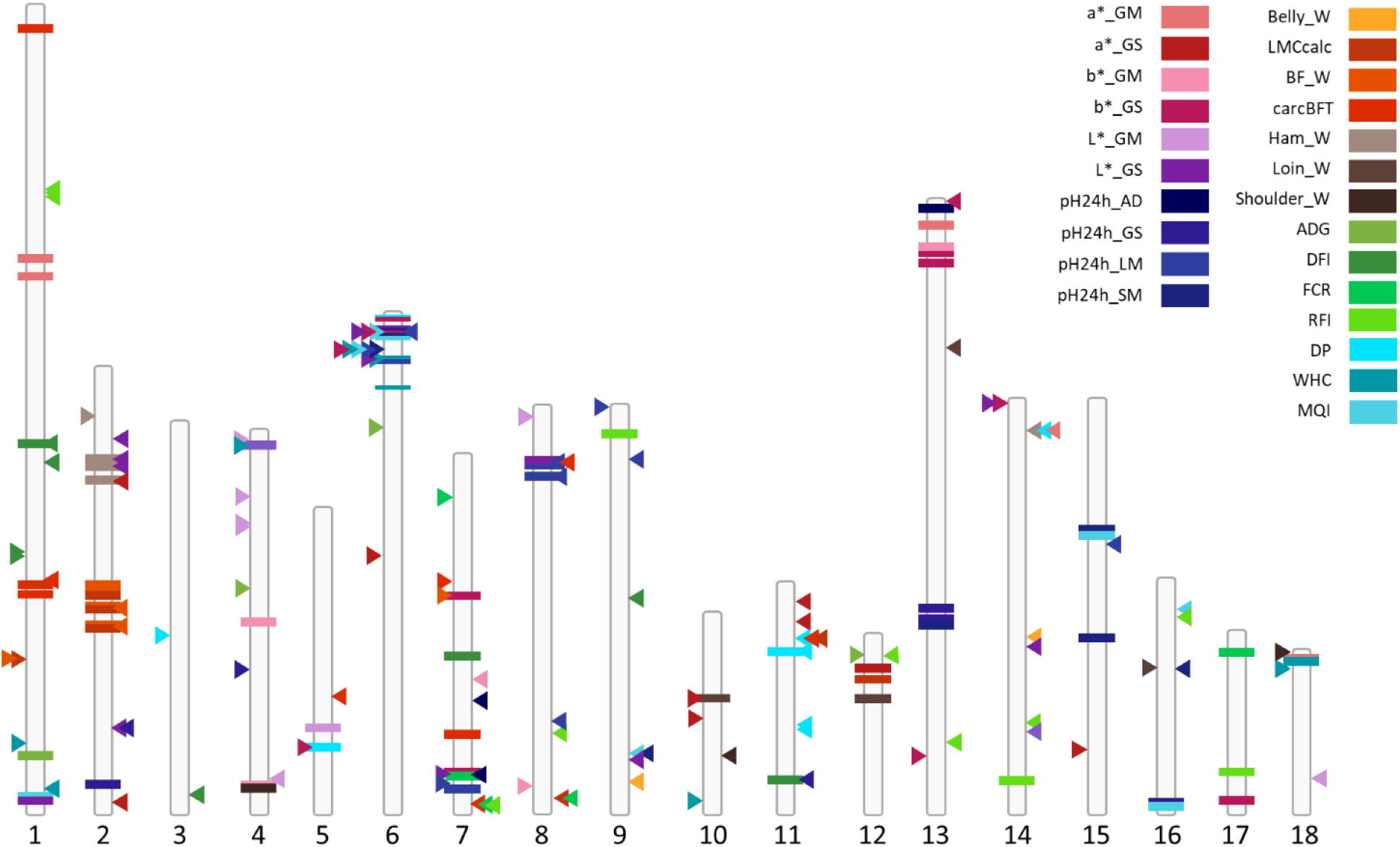
Location of all SNP-QTL identified on the 18 autosomes from the Global-GWAS, LRFI-GWAS and HRFI-GWAS. The SNP-QTL corresponding to Global-GWAS are represented by horizontal bars, LRFI-GWAS by arrows to the right of the chromosomes and HRFI-GWAS by arrows to the left of the chromosomes. Each color represents one of the 24 traits *LRFI*: low RFI line, *HRFI*: high RFI line *DFI:* daily feed intake; *ADG:* average daily gain; *FCR:* feed conversion ratio; *RFI:* residual feed intake; *carcBFT:* backfat thickness measured on carcass; *a*_GM:* a* measured on the *gluteus medius* muscle; *a*_GS:* a* measured on the *gluteus superficialis* muscle; *b*_GM:* b* measured on the *gluteus medius* muscle; *b*_GS:* b* measured on the *gluteus superficialis* muscle; *L*_GM:* L* measured on the *gluteus medius* muscle; *L*_GS:* L* measured on the *gluteus superficialis* muscle; *pH24h_AD:* pH 24h after slaughter measured on the adductor femoris muscle; *pH24h_GS:* pH 24h after slaughter measured on the *gluteus superficialis* muscle; *pH24h_LM:* pH 24h after slaughter measured on the *longissimus dorsi* muscle; *pH24h_SM:* pH 24h after slaughter measured on the *semimembranosus* muscle; *WHC:* water holding capacity of the *gluteus superficialis* muscle; *MQI:* meat quality index; *LMCcalc:* lean meat content of the carcass; *DP:* carcass dressing percentage; *Belly_W:* belly weight; *BF_W:* backfat weight; *Ham_W:* ham weight; *Loin_W:* loin weight; *Shoulder_W:* shoulder weight

To assess whether the identified QTL regions were identical and shared in the two lines, complementary GWAS analyses were performed per line, using either the set of individuals from G1 to G9 of the HRFI line or the set of individuals from G1 to G9 of the LRFI line. For analyses performed by line, the number of regions detected for a trait could differ between lines. For instance, more loci were detected in the HRFI line for b*_GS, L*_GM and WHC, whilst more regions were detected in the LRFI line for RFI, carcBFT, DP, pH24h_LM and pH24h_AD. In the HRFI line, 46 QTL were identified in 37 regions, and in the LRFI line, 68 QTL were identified in 61 regions. Only 3 regions overlapped in the two lines: on SSC6, a region located between 7 to 10 Mb affected pH24h_LM in LRFI and L*_GS, b*_GS, and MQI in HRFI, on SSC7, a region from 107 to 109 Mb affected L*_GS in HRFI and pH24h_AD in LRFI, and on SSC12, a region located between 7 to 9 Mb affected ADG in HRFI and RFI in LRFI. The two first regions affected highly correlated traits related to meat quality, but the last region affected uncorrelated traits.

Fifteen regions were shared between the 54 regions identified in the Global-GWAS and the 95 unique regions from the analyses per line, with only 3 regions common to the Global-GWAS and HRFI-GWAS analyses, 10 common to Global-GWAS and LRFI-GWAS, and the SSC6 and SSC7 regions described above detected in the three analyses (Figure 4a). Among these regions only 9 QTL (trait x region) were identified jointly in the Global-GWAS and in one of the Lines-GWAS (Figure 4b), and none was shared in the three analyses.

**Figure 4.**
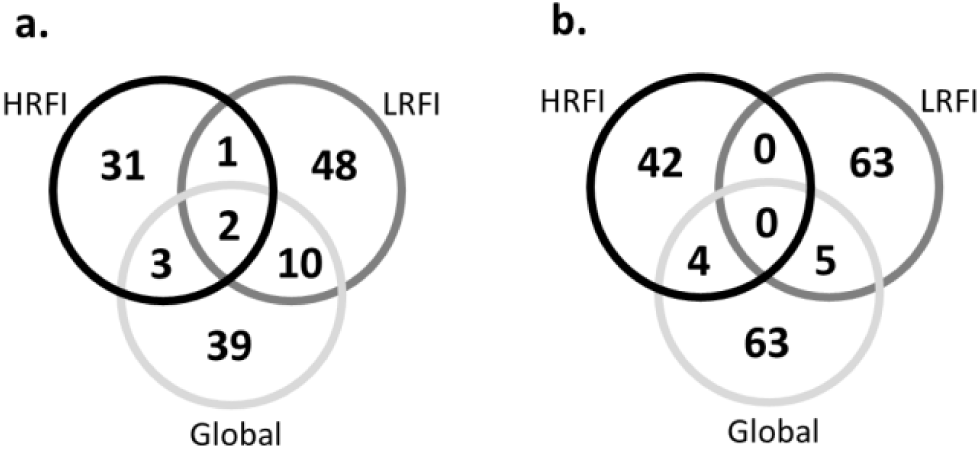
Comparison of GWAS results obtained from Global-GWAS (Global), HRFI-GWAS (HRFI) and LRFI-GWAS (LRFI). Comparison of the number of identical regions (a) and (b) comparison of the number of identical QTL (trait x region)

Very few QTL were thus common to the three GWAS (Figure 3). To assess whether a SNP-QTL significant in one analysis reached significance or suggestive thresholds in the other analyses, their *p*-values were compared. First, comparing the Lines-GWAS (Figure 5a), SNP-QTL detected *via* HRFI-GWAS had −log_10_(*p*-values) generally lower than 1 in the LRFI-GWAS, and none reached the suggestive threshold of 3. Similar results were obtained comparing SNP-QTL of the LRFI-GWAS to their *p*-values with the HRFI-GWAS. For the SNP-QTL significant in the Global-GWAS, the −log_10_(*p*-values) with the Lines-GWAS were intermediate and exceeded the suggestive threshold for several QTL. It should be noted that for these SNP-QTL, when the −log_10_(*p*-values) was suggestive in one line, it was lower in the other line.

**Figure 5.**
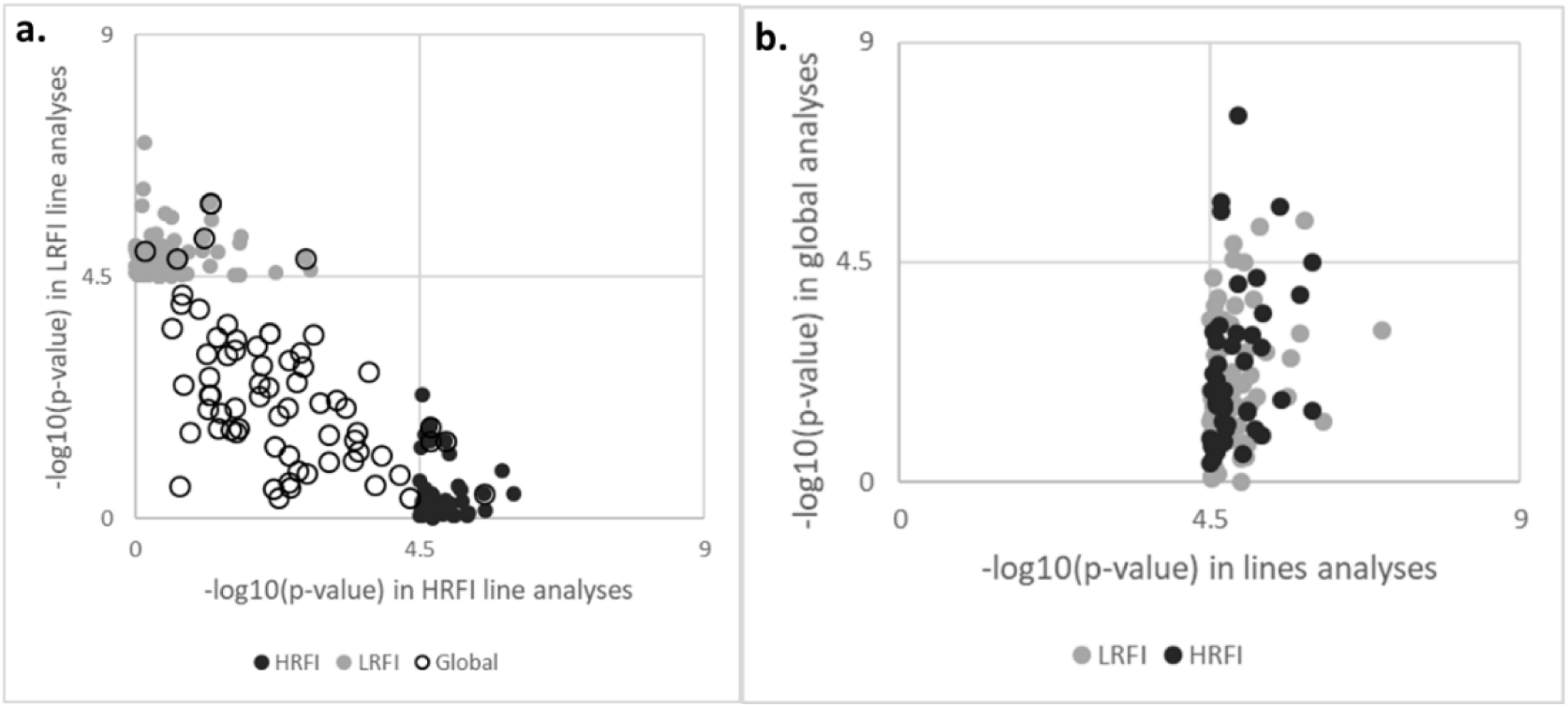
Plot of the −log_10_(*p*-value) of the SNP-QTL. The −log_10_(*p*-value) are obtained in first case with the two lines analyses for all SNP-QTL detected for the lines or the global analyses (a), and in second case obtained with the global analysis for SNP-QTL detected with the GWAS performed per line (b).

In addition, for the SNP-QTL corresponding to the QTL detected in the line analyses (HRFI-GWAS and LRFI-GWAS), the −log_10_(*p*-values) obtained in the Global-GWAS were also low (Figure 5b), with three-quarters (74.5%) of the SNP-QTL having −log_10_(*p*-values) lower than 3.

### Change of allele frequencies over the generations

The allele frequencies of the SNP-QTL detected either in Global-GWAS or in Lines-GWAS were evaluated in G1 to G9 to reflect the informativeness of these GWAS (called G9 hereafter) and in G1. When the SNP-QTL was detected in the Global-GWAS, all response animals were used to compute the frequencies; for SNP-QTL from the Lines-GWAS only the animals of the significant analysis (HRFI-GWAS or LRFI-GWAS) were used. The resulting frequency histograms are shown in Figure 6. With G1 only, 87% of the SNP-QTL of the Global-GWAS had an allele frequency between 0.2 and 0.5, with half of them between 0.4 and 0.5. In addition, 28% of SNP-QTL of the Lines-GWAS have a frequency <0.2 in G1 (Figure 6a), so the distribution of the allele frequencies of the SNP-QTL between low (<0.2) and medium was significantly different between the types of analyses (P < 0.02 for a Chi^2^ with 1 df). This difference in the distribution of the SNP-QTL allele frequencies between the two types of analyses was preserved in G9 (Figure 6b, P < 0.005): 16% of the SNP-QTL of the Global-GWAS had a frequency <0.2, compared to 35% for the SNP-QTL of the Lines-GWAS. In addition, 9% of the SNP-QTL of the Lines-GWAS had a frequency >0.6, no marker reached this frequency among SNP-QTL of the Global-GWAS.

**Figure 6.**
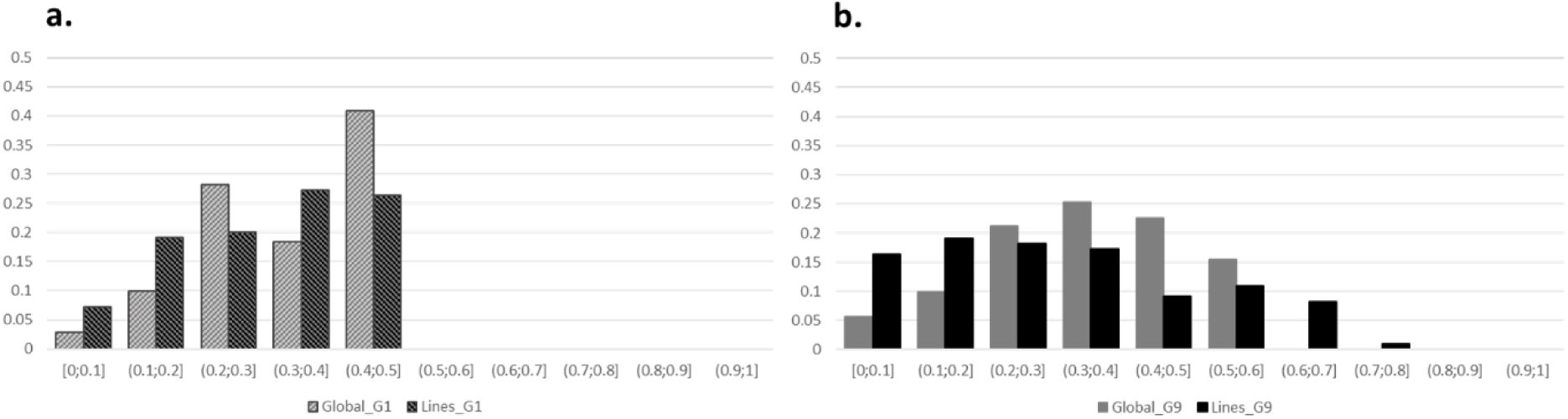
Distribution of SNP-QTL allele frequencies of Global-GWAS (in grey) and Lines-GWAS (in black). Distribution representing individuals from the line of the significant analysis (a) in G1 generation (G1 individuals only) and (b) in G9 generation (G1 to G9 individuals).

In addition to the estimation of the global allelic frequencies, we controlled if in each line the detected SNP-QTL evolved differently according to the type of analysis. First, the differences of allele frequency differences between the HRFI and LRFI lines were estimated in the G1 generation (at the beginning of the selection) (Figure 7). Regardless the analysis in which the SNP-QTL was detected, more than 65% of the SNP-QTL had low line frequency differences (<0.1) and less than 10% of the SNP-QTL had a line frequency difference >0.2. These SNP-QTL were not particularly found in one or the other type of analysis, in both types of analyses.

**Figure 7.**
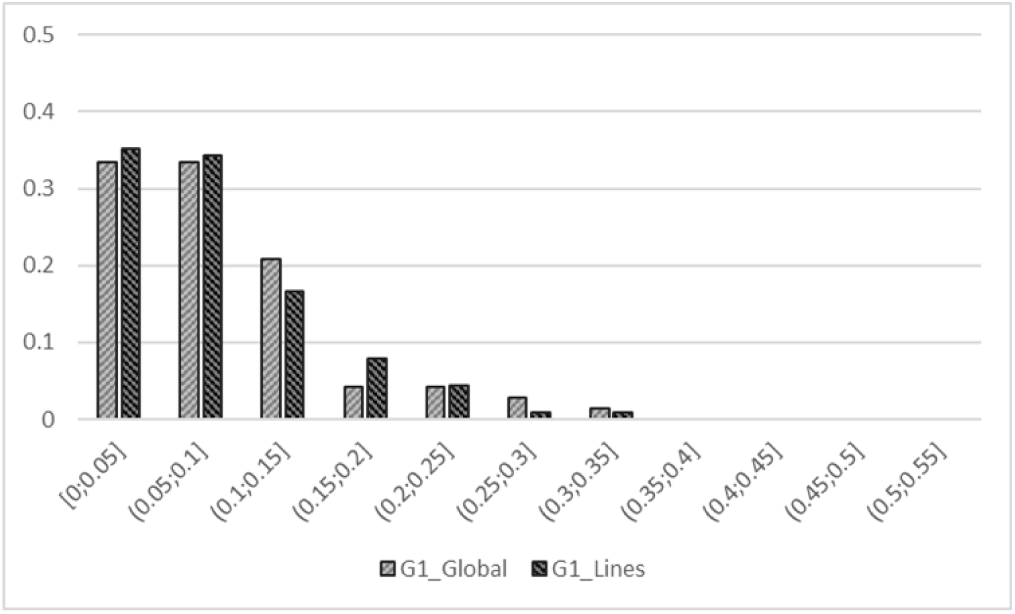
Distribution of allele frequency differences between the lines. The allele frequency differences are the absolute values between lines for SNP-QTL resulting from the Global-GWAS and Lines-GWAS in G1.

To better describe the changes of allele frequency over the generations, frequencies of SNP-QTL from Global-GWAS and Lines-GWAS were then successively estimated in each line by adding data from the next generation to the previous generations: G1 allele frequencies were obtained from G1 individuals alone, G2 allele frequencies were obtained from G1 and G2 individuals etc. Using the 9 resulting frequencies computed in each line, a linear regression of the generation number on the allele frequencies was applied within line. The comparison between lines of the regression coefficients of the allelic frequencies highlighted 4 distinct cases (Figure 8): (1) markers whose frequencies did not change with line selection (slope did not differ from zero Wald test, 5.9%), (2) markers co-selected in the two lines (slopes differed from zero and had identical sign: 22.6%), (3) markers selected in opposite directions in the lines (slopes differed from zero with different signs: 40.3%), and (4) markers whose frequencies changed only in one line (slope different from zero in one line only, 16.7% in LRFI, 14.5% in HRFI). Again no difference in the mean allele frequency evolution was observed for SNP-QTL detected in one or the other type of analysis whatever the situation (*p*-value=0.87 (No-evolution), 0.73 (Co-evolution), 0.50 (Opposite-evolution and 0.70 (One_Line_evolution) for Student T test on the values of the slopes).

**Figure 8.**
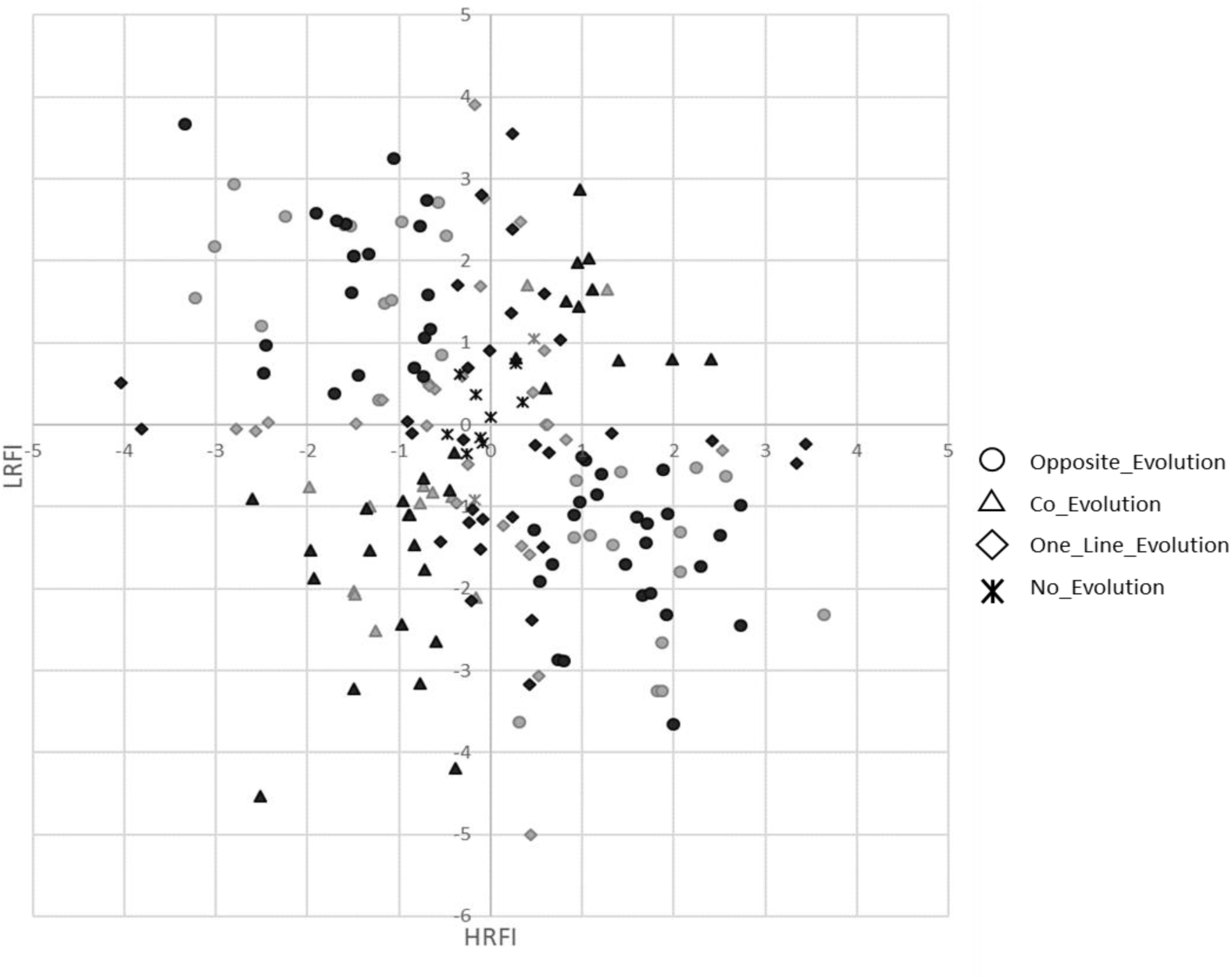
Slopes of the linear regression equations of the allele frequencies on the 9 generations. Slopes were calculated in each line, for all SNP-QTL identified with Global-GWAS (in grey) and Lines-GWAS (in black). Four situations (differentiated by different labels) were identified according to the significance of the slope (different from zero with p < 0.05 with a Wald test) in one or the two lines.

For RFI in the two lines, 9 out of the 12 detected QTL corresponded to regions identified with strong line frequencies differences: 3 RFI SNP-QTL showed differences in allelic frequency between lines higher than 0.2 in G1. The other 6 RFI SNP-QTL showed large changes of allelic frequency (regression slope > 0.024/generation). To summarize the changes of SNP-QTL allele frequencies for each trait, an average evolution score between G1 and G9 was computed using the estimated evolution scores of the different SNP-QTL detected for each trait. These averages were between 0.11 (LMCcalc) and 0.24 (RFI). A correlation coefficient of 0.66 was then estimated between the genetic line differences in G9 computed previously for the 24 different traits [27] and these averages (Figure 9).

**Figure 9.**
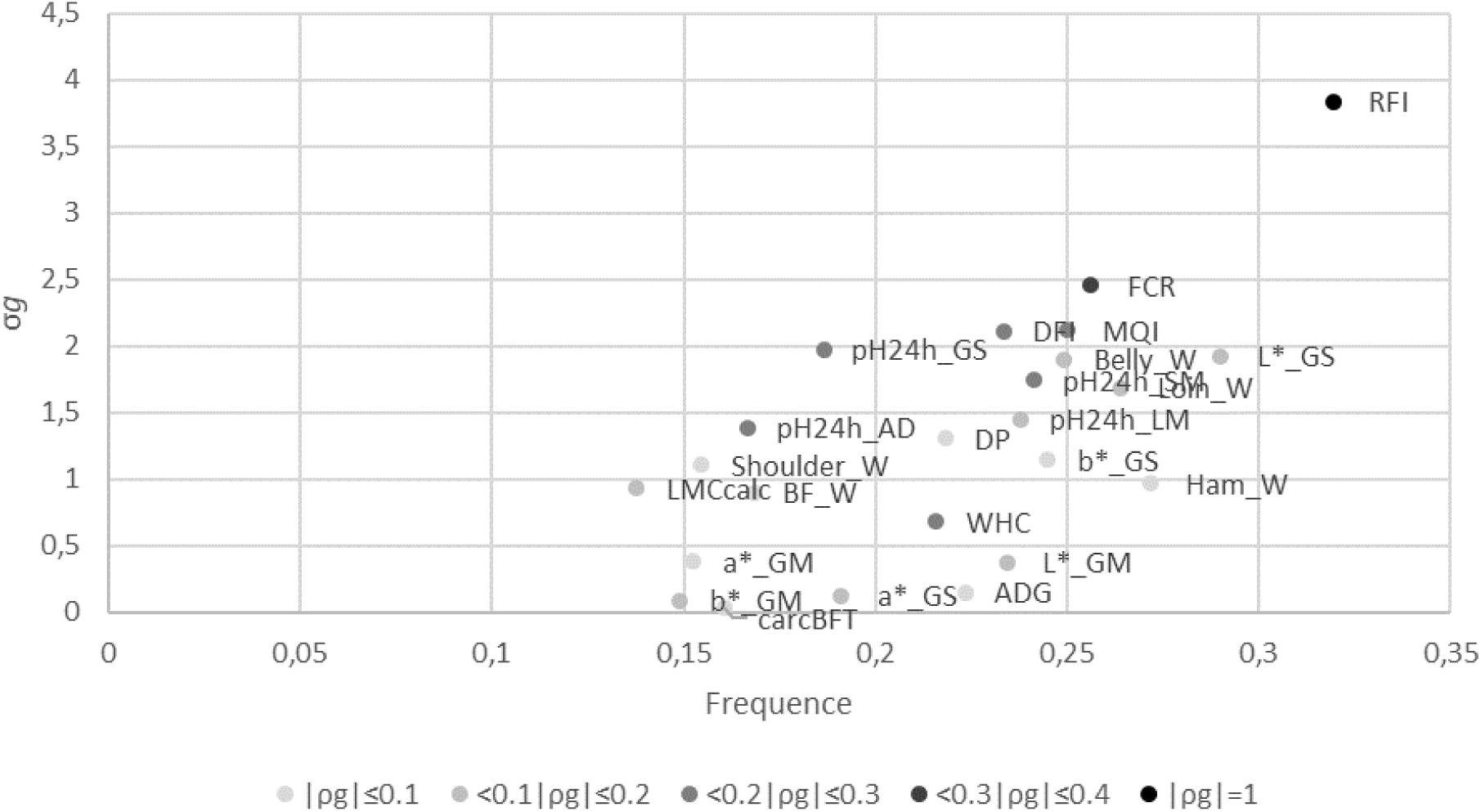
Genetic differences in G9 between the two lines. The genetic differences were expressed in genetic standard deviation of the trait (σ_g_) as a function of the average evolution of allelic frequencies in the QTL regions of the trait between the two lines. The magnitude of the genetic correlation between each trait and RFI is indicated with a grey gradient. *DFI:* daily feed intake; *ADG:* average daily gain; *FCR:* feed conversion ratio; *RFI:* residual feed intake; *carcBFT:* backfat thickness measured on carcass; *a*_GM:* a* measured on the *gluteus medius* muscle; *a*_GS:* a* measured on the *gluteus superficialis* muscle; *b*_GM:* b* measured on the *gluteus medius* muscle; *b*_GS:* b* measured on the *gluteus superficialis* muscle; *L*_GM:* L* measured on the *gluteus medius* muscle; *L*_GS:* L* measured on the *gluteus superficialis* muscle; *pH24h_AD:* pH 24h after slaughter measured on the adductor femoris muscle; *pH24h_GS:* pH 24h after slaughter measured on the *gluteus superficialis* muscle; *pH24h_LM:* pH 24h after slaughter measured on the *longissimus dorsi* muscle; *pH24h_SM:* pH 24h after slaughter measured on the *semimembranosus* muscle; *WHC:* water holding capacity of the *gluteus superficialis* muscle; *MQI:* meat quality index; *LMCcalc:* lean meat content of the carcass; *DP:* carcass dressing percentage; *Belly_W:* belly weight; *BF_W:* backfat weight; *Ham_W:* ham weight; *Loin_W:* loin weight; *Shoulder_W:* shoulder weight

## Discussion

The objective of this study was to identify QTL affecting RFI and production traits in pig lines divergently selected for RFI and to understand if the traits had different genetic backgrounds between the lines. By optimizing the genotyping to reach a good power to detect QTL in the full design and in the two lines separately, QTL were detected for all traits and hypotheses about the trait genetic background in the two lines can be formulated.

### Using average parental genotypes to detect QTL

While the use of SNPs chips now enables the genotyping of an individual at a reasonable cost, the genotyping of a design comprising several thousand individuals represents nevertheless a significant investment. In each generation of our design, at least two parities were produced, one aiming at selecting future breeders, and one to control the responses to the selection on feed consumption, growth and meat quality traits via measurements at the slaughterhouse. After 9 generations of selection, around 2,500 “response animals” had phenotypes. These individuals have the advantage of having individual records for unmeasured traits in breeders (post-mortem measurements). To optimize the costs, we genotyped all 1,632 breeders with MD SNPs chips to exhaustively survey the segregating alleles in the design. In addition, the 32 main contributors to the design were chosen from the G0 sires and dams to be genotyped using the HD SNPs chip, and an imputation step was carried out to have HD genotypes for all breeding individuals. The strong pedigree relationships in the design enabled a very good quality of HD imputation, as they help to better detect long haplotypes used to infer missing SNPs [28]. A second step was carried out, so that each response non-genotyped animal could have a genotype. These non-genotyped animal imputation have been used in cattle [29] as part of genomic evaluations to increase the size of the reference populations. In cattle, the most common situation is to determine by imputation the genotypes of dams of bulls, knowing the genotypes of the maternal grandsire, one (or more) offspring and the sires with which they were mated [30]. In such cases, the strategy takes advantage of the family information (Mendelian rule of allele transmission) and combined with allele frequencies and LD between markers at the population level. In our case, at each generation n, all response animals had both parents genotyped at generation n-1. Given these trio structures, an expected genotype at each position could be deduced from the genotypes of the parents using simple segregation rules: since the genotypes were coded as an allelic dosage for one reference allele, the genotype expectation for each offspring was simply the average of the genotypes of its two parents. As a result, 2,426 animals with genotypes (predicted) and phenotypes were available for subsequent GWAS analyses.

### Understanding the differences of detected regions between analyses

The regions detected with each type of analysis (Global- or Lines-GWAS) were very different and only 9 QTL out of 177 were shared between Global-GWAS and Lines-GWAS. The SNP-QTL detected with the Global-GWAS were far from reaching the threshold of significance in the Lines-GWAS. Similarly, most SNP-QTL detected with the Lines-GWAS were far from reaching the threshold of significance in the Global-GWAS. Although the number of individuals included in the Global-GWAS was twice higher than in the line analyses, the addition of individuals belonging to the other line seemed to have reduced the power of detection of QTL segregating in the first line. The SNP-QTL detected in the Global-GWAS or Lines-GWAS differed for their allelic frequencies in G1. This difference remained at the whole line level (G9): more SNPs with low allele frequencies were identified with the Lines-GWAS. The pedigree kinship matrix was used in the GWAS model to correct for the strong genomic structure of the population. If successful to control the type-I error of the analyses, this classical approach also limits the power of detection of QTL in highly differentiated regions between lines, as their link with the trait variability would be absorbed into the additive genetic component of the model. The Global-GWAS thus essentially allow the detection of regions segregating at intermediate frequencies in both lines. As an alternative, the analyses carried out by line allow detecting regions that got close to fixation with selection in one of the lines. From these results, it seems that the power of detection related to allele frequencies in each line is the main difference between QTL-SNPs detected with the Lines-GWAS and Global-GWAS. Given the power of the design, it is thus likely that the biological pathways involved in RFI variability in the two lines are similar, but with different contributions to the trait in each line, contrary to some previous hypotheses [10, 27].

### Comparison with published regions

Among the 12 QTL detected for RFI, three regions are detected close to RFI QTL already published. The region on SSC14 at 130-131 Mb is close to the region described by Duy N. Do et al. [31] who proposed G-protein-coupled receptor kinase 5 (*GRK5*) (129,114,449-129,343,412) as a candidate gene. Wang & al. [32] reported that a GRK5 deficiency led to insulin resistance and hepatic steatosis, and to decreases diet-induced obesity and adipogenesis in mice. In position 131,181,710-131,579,703 *FGFR2* (fibroblast growth factor receptor 2) could also be an interesting candidate. All four FGF receptors and several FGF ligands are present in the intestine and are key players in controlling cell proliferation, differentiation, epithelial cell restitution, and stem cell maintenance. *FGFR2* is expressed in the human ileum and throughout adult mouse intestine [33]. The second region closest to published RFI QTL is the 184-486 Mb interval on SSC13 near QTL reported by Bai et al. [34] and Duy N. Do et al. [31]. In this region *TMPRSS15* (transmembrane serine protease 15) is an interesting candidate gene. This gene encodes an intestinal enzyme responsible for initiating activation of pancreatic proteolytic proenzymes. It catalyzes the conversion of trypsinogen to trypsin, which in turn activates other proenzymes including chymotrypsinogen procarboxypeptidases and proelastases. *TMPRSS15* has been associated to Enterokinase Deficiency, a life-threatening intestinal malabsorption disorder characterized by diarrhea and failure to thrive [35]. On SSC17 two RFI QTL have been published by Duy N. Do et al. [31] close to the *SOGA1* gene (suppressor of glucose, autophagy-associated protein 1, 40,020,107-40,098,992) and by Onteru et al. [10] close to the *DOK5* gene (docking protein 5, 55,391,074-55,541,561). These two QTL surround the region we detected and could correspond to one unique QTL. In position 48,090,077-48,100,816, and in position 48,132,911-48,149,732, respectively, *PLTP* and *ZNF335* genes are additional candidate genes. In human, Coleman et al. [36] identified the region encoding ZNF335 as a susceptibility locus for the coeliac disease, a chronic immune-mediated disease triggered by the ingestion of gluten [36]. The PLTP (phospholipid transfer protein) transfers phospholipids from triglyceride-rich lipoproteins to high density lipoprotein (HDL). In addition to regulating the size of HDL particles, this protein may be involved in the cholesterol metabolism. PLTP KO mice absorb less cholesterol than WT mice, and have also deficient secretion by the intestine [37].

### Potential pleiotropic effects

The large number of traits recorded in our design and the known genetic correlations between these traits [27] enable the detection of pleiotropic regions, i.e. regions affecting multiple traits. Among the five regions detected for FCR, only the QTL located between 117 Mb and 119 Mb on SSC7 co-localized with a RFI QTL. For the other traits correlated to RFI (DFI, MQI, WHC, pH24h_AD, pH24h_GS, and pH24h_SM traits), only 3 QTL were detected within 10 Mb of the RFI QTL: a QTL at 2 Mb influencing MQI on SSC16 between 11 and 12 Mb, and two QTL on pH24h_AD at 7 Mb and 10 Mb of QTL for RFI located at 113-114 Mb on SSC14 and 107-109 Mb on SSC7, respectively. Compared to the previously published QTL regions for RFI, we identified a QTL influencing FCR in a region described by Onteru et al. [10] between 15 and 16 Mb on SSC7, a QTL for pH24h_SM in the 80 and 81Mb interval on SSC15 described by Duy N Do et al. [31], and a QTL for DFI in the region described by Y M Guo et al. [38] on SSC3 between positions 126 and 128Mb. Despite the reported correlations between these traits and RFI, among the 52 QTL detected in our study for DFI, MQI, WHC, pH24h_AD, pH24h_GS, and pH24h_SM, only seven co-located with RFI QTL identified in our study or in previously published studies.

### Changes of QTL allele frequencies and trait responses to selection

The allele frequencies of the majority of the detected regions changed between the G1 and G9 generations, with more than 70% of the regions for which SNP-QTL evolved in opposite directions or in a one line only. However, the magnitude of allelic changes of the QTL regions varied from one trait to the next, and was strongly correlated with line differences previously reported in G9 [27]. Indeed, the regions with the highest allele frequency changes were detected for RFI, which was trait used for selection. For the other traits, the higher the genetic correlation with RFI, the higher the frequency variation of the associated QTL regions. As a result, QTL affecting FCR, DFI and MQI had the highest frequency changes with generations. The responses of QTL affecting meat quality traits are consistent with the high and early responses to selection previously detected in this experimental population for these traits [5]. Altogether, our analyses underline a clear relationship between the quantitative responses to selection of the traits and changes of alleles frequencies in some QTL regions, certainly pointing out chromosomic regions that were selected during the experiment, whereas in such populations of low effective size and strong directional selection, detecting selection signatures with standard methodologies [39] can have low power due to the major effect of drift on the changes of the allele frequencies. However, recently developed new methods, based on genetic time series could provide new insights for the detection of regions under selection in small populations [40].

## Conclusions

This study aimed at characterizing the molecular architecture of RFI in two lines divergently selected for this trait. Besides efficiently detecting known and new QTL regions, the combination of GWAS carried out per line or simultaneously using all individuals allowed the identification of candidate regions of the genome under selection, which can explain the responses to selection of different traits reported before. Analyzing the allelic frequencies from G1 to G9, we concluded that the majority of the QTL regions responded to selection in a divergent way in the lines, and that the same metabolic pathways were certainly involved in both lines. Several new regions determining RFI variability were identified in this study and new candidate genes were proposed to complement the data acquired in previous published analyses.

## Declarations

### Consent for publication

Not applicable

### Availability of data and materials

The datasets used and/or analysed during the current study are available from the corresponding author on reasonable request.

### Authors’ contributions

ED performed the statistical analyses and wrote the first draft of the paper. YB and KF organized the data acquisition. ED and YL performed the imputation and quality control of the genotypic data. ED, AA, YL, HG and JR participated in the design of the study. JR and HG provided scientific supervision. All authors read and approved the final manuscript.

## Acknowledgements

The authors would like to thank (*i*) the experimental farm staff for data collection, samples management and breeding of the animals and (*ii*) both technology platforms, CRCT and Gentyane, for the genotyping.

## Additional files

**Additional file 1**

Format: additionalfile1.xlsx

Title: Number of animals used for the analyses after quality control

Description: Details of the number of aniamls before and after application of filter on the call rate (CR) were given for chips (60K, 70K and 650K SNPs chips), imputation levels (MD/HD imputation) and average genotypes calculated from the genotypes of both parents (HD predicted).

**Additional file 2**

Format: additionalfile2.xlsx

Title: Number of SNPs used for the analyses after quality control

Description: Details of the number of SNPs before and after application of filters on the call frequency (CF) and the frequency of minor allele (MAF) were given for chips (60K, 70K and 650K SNPs chips), imputation levels (MD imputation and HD imputation) and average genotypes calculated from the genotypes of both parents (HD predicted).

**Additional file 3**

Format: additionalfile3.xlsx

Title: QTL regions detected with the three groups of association studies

Description: These QTL regions were found from the full population (Global-GWAS) and from each line separately (HRFI-GWAS and LRFI-GWAS)

*DFI:* daily feed intake; *ADG:* average daily gain; *FCR:* feed conversion ratio; *RFI:* residual feed intake; *carcBFT:* backfat thickness measured on carcass; *a*_GM:* a* measured on the *gluteus medius* muscle; *a*_GS:* a* measured on the *gluteus superficialis* muscle; *b*_GM:* b* measured on the *gluteus medius* muscle; *b*_GS:* b* measured on the *gluteus superficialis* muscle; *L*_GM:* L* measured on the *gluteus medius* muscle; *L*_GS:* L* measured on the *gluteus superficialis* muscle; *pH24h_AD:* pH 24h after slaughter measured on the adductor femoris muscle; *pH24h_GS:* pH 24h after slaughter measured on the *gluteus superficialis* muscle; *pH24h_LM:* pH 24h after slaughter measured on the *longissimus dorsi* muscle; *pH24h_SM:* pH 24h after slaughter measured on the *semimembranosus* muscle; *WHC:* water holding capacity of the *gluteus superficialis* muscle; *MQI:* meat quality index; *LMCcalc:* lean meat content of the carcass; *DP:* carcass dressing percentage; *Belly_W:* belly weight; *BF_W:* backfat weight; *Ham_W:* ham weight; *Loin_W:* loin weight; *Shoulder_W:* shoulder weight

